# *Gekko gecko* as a model organism for understanding aspects of laryngeal vocal evolution

**DOI:** 10.1101/2024.05.10.593509

**Authors:** Ruth Gutjahr, Loïc Kéver, Thorin Jonsson, Daniela Talamantes Ontiveros, Boris P Chagnaud, Anthony Herrel

**Author notes:** **Corresponding author:** Ruth Gutjahr. Authors contributed equally (Ruth Gutjahr, Loïc Kéver, Boris P Chagnaud, Anthony Herrel).

## Abstract

The ability to communicate through vocalization plays a key role in the survival of animals across all vertebrate groups. While avian reptiles have received much attention relating to their stunning sound repertoire, non-avian reptiles have been wrongfully assumed to have less elaborate vocalization types and little is known about the biomechanics of sound production and their underlying neural pathways. We investigated alarm calls of *Gekko gecko* using audio and cineradiographic recordings of their alarm calls. Acoustic analysis revealed three distinct call types: a sinusoidal call type (type 1), a train-like call type, characterized by distinct pulse trains (type 3), and an intermediary type, which showed both sinusoidal and pulse train components (type 2). Kinematic analysis of cineradiographic recordings showed that laryngeal movements differ significantly between respiratory and vocal behavior: during respiration, animals repeatedly moved their jaws to partially open their mouths, which was accompanied by small glottal movements. During vocalization, the glottis was pulled back, contrasting with what has previously been reported. *In-vitro* retrograde tracing of the nerve innervating the laryngeal constrictor and dilator muscles revealed round to fusiform motoneurons in the hindbrain-spinal cord transition ipsilateral to the labeled nerve. Taken together, our observations provide insight into the alarm calls generated by *G. gecko*, the biomechanics of this sound generation and the underlying organization of motoneurons involved in the generation of vocalizations. Our observations suggest that *G. gecko* may be an excellent non-avian reptile model organism for enhancing our understanding of the evolution of vertebrate vocalization.

**Summary Statement:** Investigation of *Gekko gecko* alarm calls revealed distinct call types, during which the larynx is being pulled back by muscles innervated by motoneurons located in the hindbrain.

## Introduction

For vertebrates, the ability to communicate through vocalization is a key feature for survival for many species. The behavioral role of this communication channel has been investigated for every major vertebrate group (Amorim, 2006; Hollén and Radford, 2009; Ryan and Rand, 2001; Slabbekoorn and Smith, 2002). Signaling phenotypes are central to most animal life and variations in their features have promoted bursts of speciation (Laiolo, 2010; Streelman and Danley, 2003). Insights into the sound production apparatuses and the mechanisms by which the neuromuscular systems control sound production have been provided for diverse taxa including fish, amphibians, avian reptiles, and mammals (Suthers et al., 2016). While detailed anatomical descriptions and a few observations of laryngeal movements and vocal cords during sound production (Paulsen, 1967; Russell and Bauer, 2021) are present, the biomechanics and the neural pathways controlling vocalizations in non-avian reptiles remain poorly studied. Although vocalizations with different levels of complexity have evolved repeatedly among non-avian reptiles (Jorgewich-Cohen et al., 2022), the imprint of acoustic communication on their evolutionary success and diversity has been generally underappreciated (Russell and Bauer, 2021). Compared with other vertebrate groups, the role, control, and tuning of their communication, as well as the role acoustic communication has played in their diversity and evolutionary history remains much less well known.

Most non-avian reptiles have long been considered to be silent or to possess a poor vocal repertoire (Russell and Bauer, 2021) which probably explains why they are the only large vertebrate group in which the central and peripheral mechanisms controlling the sound features remain largely obscure. This is regrettable because it is now undisputed that vocalizations with different levels of complexity have evolved repeatedly among non-avian reptiles (Russell and Bauer, 2021). An in-depth understanding of the mechanisms controlling their sound production system(s) would be helpful to understand basic mechanisms of complex vocalizations in general.

Studies conducted on the sound production apparatus of non-avian reptiles date back to the 19^th^ century (Humboldt, 1811; Moore et al., 1991; Russell and Bauer, 2021; Vergne et al., 2009), but have focused almost exclusively on the gross morphology of the laryngotracheal complex. Among reptiles, “true” vocal cords are present only in some gekkotans even though some other lizard species and turtles have ligament-like structures that could play a similar role (Moore et al., 1991; Russell and Bauer, 2021). Henle (1839) described an impressive diversity in laryngeal morphology across the Gekkota. However, only a single study has described the laryngeal movements during distress calls in a gecko (Paulsen, 1967), limiting our understanding of the relationship between the observed morphological diversity and the diversity in sound production.

To date, it is generally accepted that non-avian reptiles generate and modulate sounds at the level of the glottis, most likely by involving constrictor and dilatator muscles acting on the cricoid and arytenoids (Colafrancesco & Gridi-Papp, 2016). This sound production apparatus is located at the cranial end of the trachea, in contrast to the syrinx of avian reptiles which is positioned at its caudal (or posterior) extremity. This anatomical feature makes the sound production apparatus of non-avian reptiles much easier to access and may thus facilitate *in vivo* analyses. Despite the fact that most vocal reptiles generate relatively simple sounds that may not be species-specific (*e.g.,* hissing), other species are known to generate several call types with species-specific features (Rohtla Jr. et al., 2019; Yu et al., 2011). In the gecko *Chondrodactylus turneri* (Vedenin, 2015), for example, the features of the courtship call are more conserved than those of the distress call, suggesting that it contains species-specific information.

We here investigate sound production aspects in the Tokay gecko (*Gekko gecko*). We chose this species, as it is highly vocal and produces several easily elicited sound types (alarm calls; Fig. 1), even in captivity (Brumm and Zollinger, 2017; Paulsen, 1967; Tang et al., 2001; Yu et al., 2011). We investigated the acoustic properties of these readily-inducible vocalizations in a laboratory context and explored the motion of jaws and the position of the larynx during respiration and vocal behavior. We further studied the innervation of the laryngeal constrictor and dilator muscles by tracing the motor neurons using an *in vitro* preparation. Combined, our data highlight the suitability of these animals for in-depth neurophysiological studies (Kennedy, 1975; Kennedy, 1981) and as a model for our understanding of vocal communication in non-avian reptiles.

**Figure 1:**
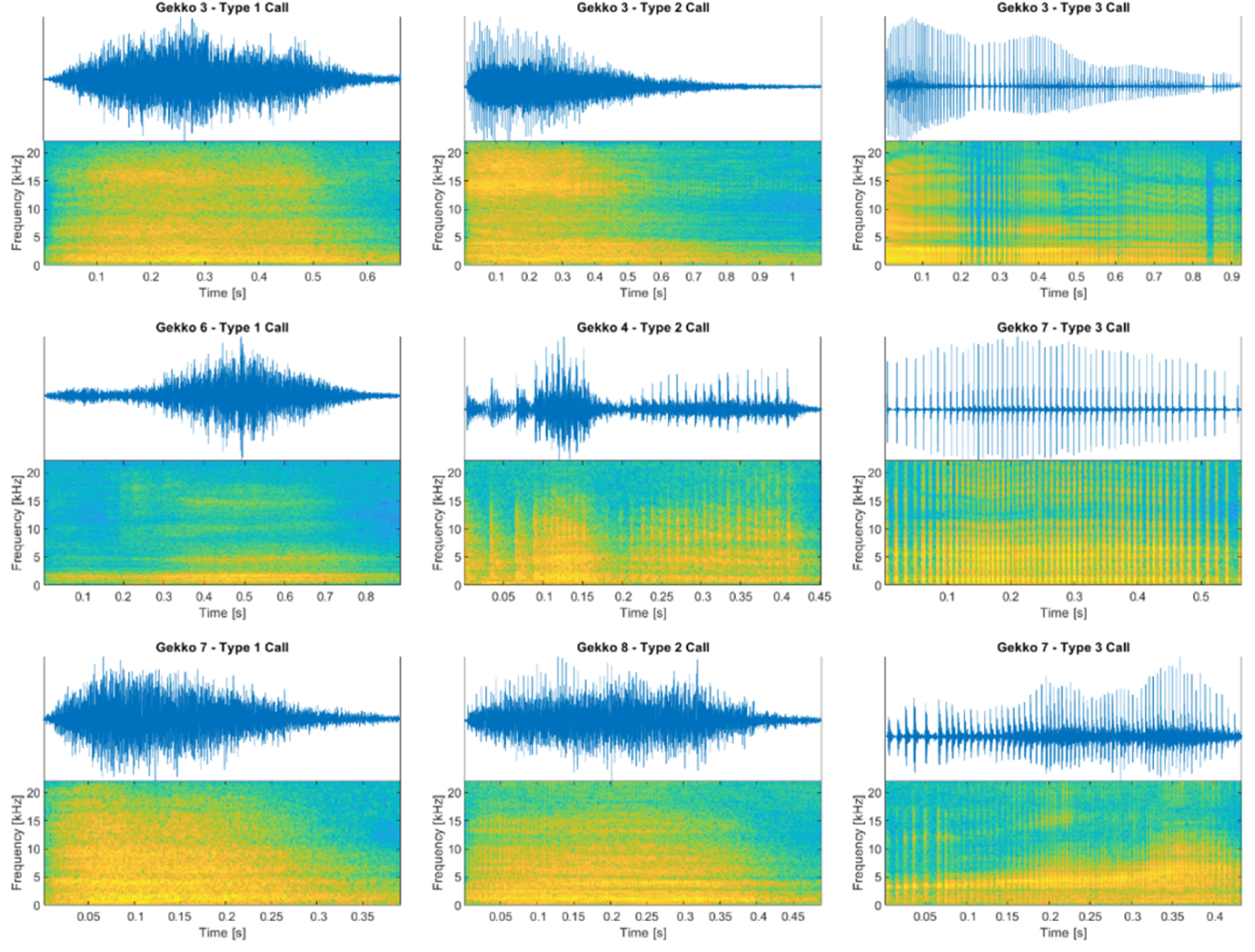
Examples of the three visually identified types of alarm call vocalizations (n = 156) produced by individual *Gekko gecko* (N = 6). Left column: Type 1 calls. Middle column: Type 2 calls. Right column: Type 3 calls. Upper part shows the amplitude waveform of the call, the lower part the matching spectrogram (see text for spectrogram details). Amplitudes are normalized for each call and spectrogram colours indicate power at the corresponding frequency and moment in time; blue: low power, yellow: high power.

## Materials and methods

### Animal care and maintenance

Animals (*Gekko gecko*, Linnaeus) of both sexes (8 males and 2 females) were acquired from the commercial trade (Paris*: La Ferme Tropicale* and Graz: *Zoo Muser*) and ranged in size from 119.0 to 148.2 mm (SVL) and 28.30 to 50.07 g in body mass. Animals were held in terraria under a 12:12 dark:light cycle in temperature-controlled rooms at 25 °C. Animals were sprayed daily with tap water and fed with live grasshoppers and crickets. Observations in Paris were conducted in accordance with the ethics guidelines of the Comité Cuvier at the Muséum national d’Histoire naturelle (MNHN) under the animal protocol #68.

### Sound recordings

For acoustic analysis, six adult *Gekko gecko* individuals were used (for details on the exact sample sizes, See Table S1). Animals were hand-held during animal maintenance procedures, during which they readily elicited vocalizations. We focused on alarm calls as a class of vocal signals as these were most easily elicited and could be analyzed both in terms of kinematics, as well as for sound signal properties. Sounds were recorded with a microphone (model 472049; MicW, Beijing, China) placed 25 cm in front of the respective animal. Microphone signals were digitized with an external sound card (Motu UltraLite mk3, Motu Inc., Cambridge, MA, USA) at 44.1 kHz sampling rate using Audacity (v. 3.2.4, audacityteam.org). A 100 Hz high-pass filter was applied to the recorded calls before sequences were saved on a PC as 16-bit wave files. Two recording sessions were performed at 25 °C for each animal, resulting in 12 recorded call sequences ranging in duration from ∼1-2 minutes.

### Acoustic analysis

Individual alarm call vocalizations were excised from the recorded call sequences, visually categorized into one of three call types (Fig. 1) and saved as separate wave files (using Audacity). This resulted in a total of 156 calls recorded from six animals (N = 6, n = 156; see supplementary table 1 for details about the number of calls per type and individual animal). All subsequent acoustic analyses were carried out using custom written code in Matlab (R2021b; The MathWorks Inc., Natick, MA, USA). To visualize the frequency composition of calls in time, spectrograms of selected recordings were calculated using Matlab’s *spectrogram* function with 4096 Fast Fourier Transform (FFT) points and a Hamming window of variable size (window size = total number of samples per signal/100; resulting in 100 time bins per signal with a frequency resolution of 10.8 Hz). To investigate the frequency composition of the gecko calls, spectral analysis was carried out on each individual vocalization by first applying a 250 Hz high-pass filter (6^th^ order Butterworth) to the acoustic signals, followed by Welch’s power spectral density (PSD) estimate with 4096 FFT points and a 256-point Hanning window. Resulting spectra for call types were averaged per individual, after which the mean over the individual averages was calculated to show the mean PSD of each call type. Averages over all calls produced by an individual were calculated as well.

#### Principal components analysis of vocalizations

For each recorded vocalization (N = 6, n = 147; nine recordings were not included in the principal components analysis because of their short duration), acoustic feature vectors in the form of 15 mel-frequency cepstral coefficients (MFCC) and corresponding delta coefficients for 20 segments (window size = total number of samples per signal/20) were calculated using *melfcc* (see Clink and Klinck, 2021; Ellis, 2012 for a detailed description of the method). To cover the hearing range of G. gecko, the minimum and maximum band edges of the mel filter were set to 250 and 5000 Hz, respectively (Manley et al., 1999; Manley et al., 2014). The first MFCCs (also denoted as 0^th^ coefficient or MFCC 0) and its deltas were omitted from the analysis as these relate to the overall energy of the signals only (Clink and Klinck, 2021; Clink et al., 2019). Subsequently, principal component analysis (PCA) was performed on the resulting vocalization feature vectors to calculate the first three component coefficients and scores.

### High speed cineradiography

Prior to videography, two adult males (N = 2, for details on the sample sizes, see Table S2) were implanted with small metal markers to help quantify movements of the jaws and the larynx. Markers were placed subcutaneously at the anterior and posterior extremities of the upper and lower jaws using a hypodermic needle. One marker was placed next to the laryngeal cartilages allowing the tracking of movements of the anterior-most part of the respiratory system. All markers were inserted under light anesthesia achieved by intramuscular injection into the hind limbs of a mixture of ketamine (80 mg/kg) and medetomidine (115 µg/kg; see Barrillot et al., 2018; Chai et al., 2009). After implantation anesthesia was reversed by an intramuscular injection of atipamezole (115 µg/kg).

X-ray videos were recorded at 200 to 400 fps at a resolution of 1280*800 pixels. Two Phantom Miro cameras (model R311; AMETEK, Berwin, PA, USA) mounted on the image intensifiers of a custom-designed biplanar X-ray system (RST Medical; https://rstmedical.nl/) were used to record a side and a top view of the geckos (see suppl. videos 1-4). The cameras were mounted on Philips Imagica 38 cm imaging systems (Koninklijke Philips N.V., Amsterdam, Netherlands) mounted on Alp Lift L lift-trolley systems (Alp Lift B.V., Utrecht, Netherlands). X-rays were generated at 57 kV at 80 mA and 60 kV at 80 mA using Philips Super 50CP X-ray generators and Philips SRM 1550 ROT350 X-ray tubes also mounted on Alplift L lift-trolley systems. Sequences were recorded for six seconds simultaneously in lateral and dorso-ventral views. Geckos were positioned inside a transparent acrylic tube (diameter: 5 cm) and were incited to vocalize by approaching them with a piece of black foam. Recordings were performed in an environmentally controlled room set at 25 °C. During the X-ray video recordings, sounds were acquired at a distance of 1.5 m with a laptop sound card and internal microphone using the audacity software.

### X-ray video analysis

For each individual (N = 2, for details on the sample sizes, see Table S2), five vocalization and two breathing sequences were analyzed. In each six-second sequence multiple breathing or vocalization events were included. Both camera views were analyzed separately due to a problem with camera synchronization during recording. Markers were tracked using the ProAnalyst software (Xcitex Inc., Woburn, MA, USA) and coordinates were exported to Microsoft Excel, smoothed using a low pass Butterworth filter and scaled. Based on the marker coordinates the distance between the two anterior jaw markers was calculated and is referred to hereafter as gape distance. We also calculated the distance between the laryngeal marker and the anterior marker on the lower jaw in dorsal view (i.e., a measure of antero-posterior displacement of the glottis relative to the lower jaw) and the distance between the posterior marker on the upper jaw and the laryngeal marker in lateral view (i.e., a measure of the dorso-ventral displacement of the glottis relative to the upper jaw). From the kinematic profiles we extracted the total movement of the glottis and the duration of the movement cycle for each vocalization and breathing event recorded (N = 2 animals, Gecko 1 lateral view: *n*_breathing_ = 5; *n*_vocalization_ = 14; Gecko 2 lateral view: *n*_breathing_ = 6; *n*_vocalization_ = 6; Gecko 1 dorsal view: *n*_breathing_ = 8; *n*_vocalization_ = 8; Gecko 2 dorsal view: *n*_breathing_ = 5; *n*_vocalization_ = 17). To test for differences in laryngeal movement between breathing and vocalization, we first Log_10_-transformed the absolute displacements and then ran a general linear model testing for differences between the two behaviours. We then also ran a similar analysis on the Log_10_-transformed duration of the movement cycle. Finally, we ran simple linear regressions between the jaw gape profile and the laryngeal movement profile to explore whether jaw and laryngeal movements were coordinated. From these regressions we extracted the R^2^ value as a measure of the strength of association between the jaw and laryngeal movement. All analyses were run in IBM SPSS statistics V.29.

### Neuronal tracing

We used 4 male animals (N = 4, for details on the sample size, see Table S2) (119.0 - 127.7 mm SVL, 28.30 - 34.72 g) for *in-vitro* nerve backfills. For these tracings, animals were deeply anesthetized using ketamine (150 mg/kg, Richter Pharma, Wels, Austria), perfused transcardially using a carbogenized, ice-cold snake ringer solution (Bothe et al., 2018) and were subsequently decapitated, leaving the target nerves intact. We removed excess tissue to gain access to the glossopharyngeal nerve, which served as the target nerve for our retrograde tracings. Ringer solution was removed and the nerve was severed using microscissors. We used tracer-coated insect needles to place a small BDA-crystal (biotinylated dextran amine, 3kDa; Invitrogen, Eugene, OR, USA) onto the proximal end of the severed nerve and incubated the preparation for 10 min. After this period, excess dye was rinsed off using fresh, ice-cold ringer solution. For the incubation period, the tissue was placed into a 1 l glass beaker filled with ringer, which was continuously carbogenized. The total tracer incubation time was 48-72 hours at 4 °C and ringer solution was exchanged every 12 hours.

After incubation, the tissue was immersion-fixed in 4% paraformaldehyde (PFA) in 0.1 M PB overnight and washed in 0.1 M PB before removing the brain from the skull. Brains were embedded in 2% agarose (ROTH, Karlsruhe, Germany) in 0.1 M PB and transversely sectioned at 100 µm on a vibratome (7000smz-2; Campden Instruments, Loughborough, UK). Sections were mounted onto chrome alum-gelatine coated slides and left to dry overnight. Next, the sections were washed in 0.1 M PB for five minutes, permeabilized with 0.5% Triton X-100 (Tx-100) in 0.1 M PB for 10 minutes and developed with a streptavidin-solution (1:500 in 0.5% Tx-100 in 0.1 M PB, Life Technologies, Carlsbad, CA, USA). The tissue was washed three times in 0.1 M PB and once in 0.025 M PB. Sections were mounted using VECTASHIELD Antifade Mounting Medium with 4′,6-diamidino-2-phenylindole (DAPI, Vector Laboratories, Burlingame, CA, USA).

### Imaging and quantification

Slices were inspected and imaged on an epifluorescence microscope (Thorlabs Cerna Confocal 208 CM712, Newton, NJ, USA and ZEISS Axio Imager 2, ZEISS, Jena, Germany). For better visualization of the labeled neurons we adjusted brightness and contrast as necessary. Imaged slices were adjusted and analyzed using Fiji (v. 1.52, Schindelin et al., 2012). We determined minimal (min.) and maximal (max.) soma diameter and the surface area of the traced neurons. Analyses were performed using a custom script in python 3 (v. 8.3), building on the libraries ‘*pandas’* (v. 2.0.1(McKinney, 2010)) and ‘*seaborn’* (v. 0.11.2, (Waskom, 2021)). We calculated the minimum, maximum, mean and standard error of the mean (SEM) of the neuroanatomical measurements. Histograms of min., max. soma diameter and surface area of the neurons were plotted, using the following bin sizes: min. and max. diameter = 4 µm; surface area = 100 µm².

## Results

### Sound quantification

A total of 156 vocalizations were obtained from six individuals. Geckos did not emit their characteristic “to-keh” advertisement call under hand-held conditions. An initial visual analysis of the waveforms of the recorded call sequences suggested that the geckos produced three different types of vocalizations: a sine-like call (type 1 call; Fig. 1 left column), a train of irregularly spaced individual pulses (type 3 call, Fig. 1 right column), and a mixture of both, where pulse trains are visible within sinusoidal events (type 2 call, Fig. 1 middle column). Of the 156 calls (N = 6), 54 vocalizations were of type 1, 34 of type 2, and 68 of type 3 (see Table S1 for a distribution of call types by individual). Sounds were characterized by a relatively broad frequency range (∼85% of all power between 500 Hz and 5 kHz), with a mean peak frequency around 1 kHz for all call types and were similar in frequency content across call types and individuals (Fig. 2 and Fig. S1).

**Figure 2:**
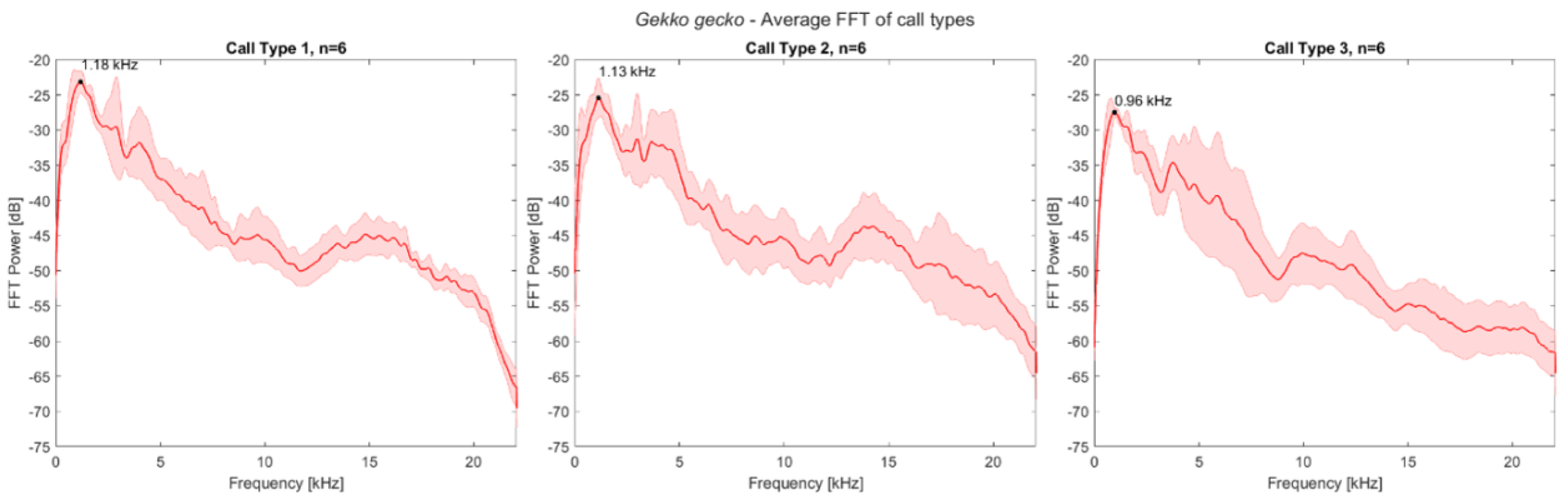
Mean frequency spectra of the three *Gekko gecko* alarm call types. Solid line shows the average (N = 6 animals, n = 156 calls), shaded areas ± 1 standard deviation. Black dot marks the frequency with the greatest amplitude.

To test whether the visually identified vocalization types do indeed represent distinct types of alarm calls, a principal component analysis of the signals’ mel-frequency cepstral coefficients (MFCC) was carried out. Cepstral coefficients are a form of acoustic feature vectors and can be extracted from the signal as shape descriptors of the spectral envelope (Clink and Klinck, 2021). Here, the non-linear mel scale was chosen for frequency mapping instead of linearly spaced frequency bands because most terrestrial vertebrates do not perceive sound frequency in a linear manner (Deecke and Janik, 2006; Manley et al., 1999). For each vocalization (n=147, N = 6; nine recordings were excluded due to short duration), 15 MFC and delta coefficients were calculated and PCA was performed on those feature vectors. The first three component scores for all vocalizations and their corresponding call types were then plotted in 2D and 3D to investigate if the visually determined call types correspond to observable clusters in the principal component space (Fig. 3 and Fig. S2). Two separate clusters delineating type 1 and type 3 calls were seen with type 2 calls broadly overlapping call types 1 and 3. Thus, we concluded that type 1 and type 3 calls are indeed different call types, and that type 2 calls represent an intermediate call type sharing properties of both type 1 and type 3 vocalizations.

**Figure 3:**
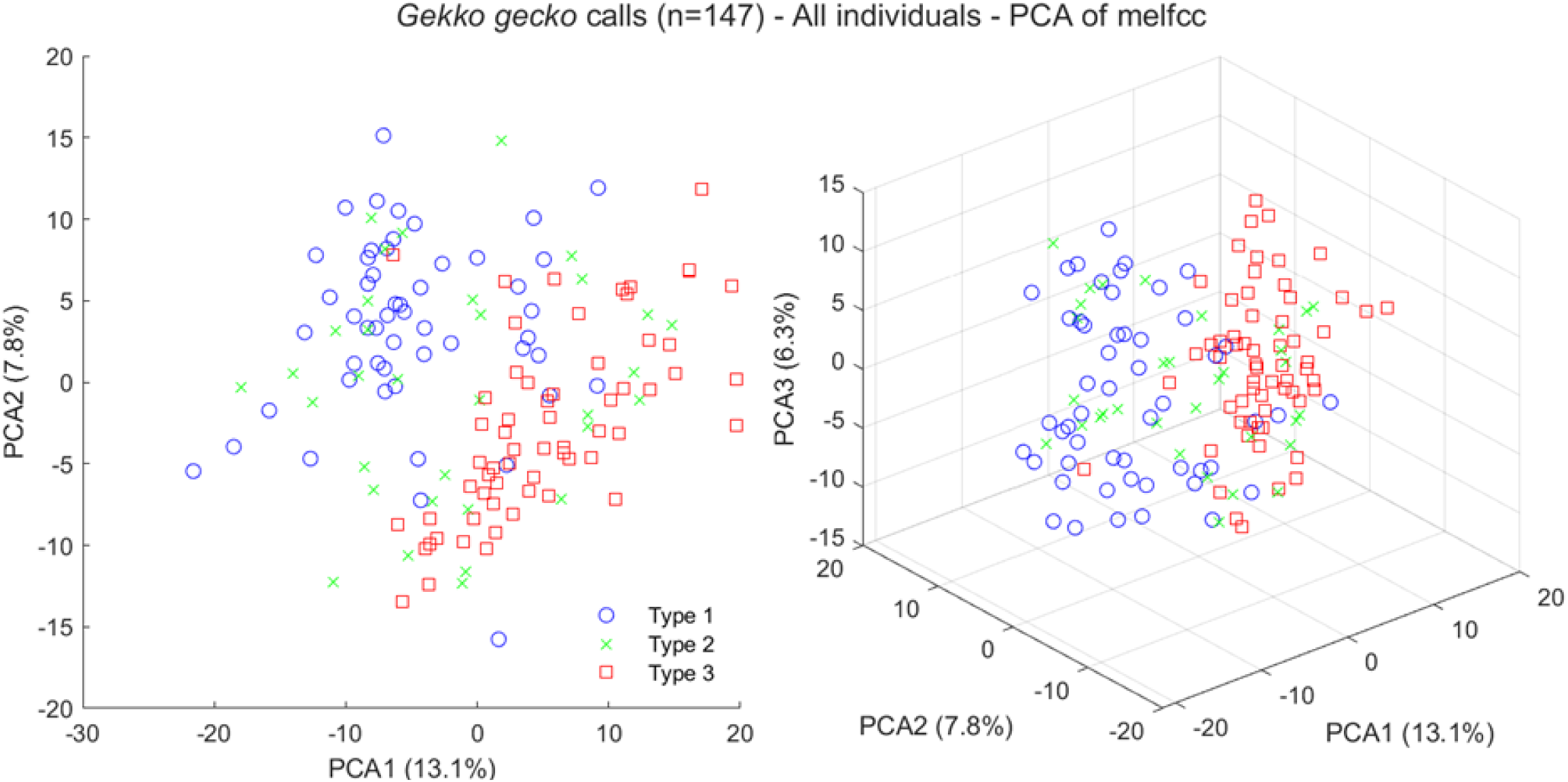
Results of PCA of MFCCs of alarm calls (n = 147) produced by *Gekko gecko* individuals (N = 6). Left: 2D plot of the first two principal components, explaining 20.9% of the total variance. Blue circles, green crosses and red squares represent the call types (Type 1, 2 and 3, respectively) assigned to the vocalizations after visual inspection of the wave forms. Right: 3D plot of the first three principal components of the same data, explaining 27.2% of the total variance.

### Kinematic analysis

#### Breathing

During breathing, the mouth opens slightly in a repeated pattern roughly associated with laryngeal movements (Figs. 4, 5). More specifically, as the mouth opens slightly, the larynx is pulled rostrad, closer to the tip of the lower jaw. However, most of the movements occur in the vertical plane with the larynx being pulled ventrally relative to the largely stationary lower and upper jaws (Supplementary videos S1-2 and see Fig. S3 for schematic positioning of the markers). Displacements of the jaws are very small (< 1 mm, Fig. 4, 5). Although displacements of the larynx in the horizontal plane are also very small (typically less than 1 mm, mean = 0.06 ± 0.03 cm), displacements in the vertical plane during breathing are larger and can be up to 3 mm (mean = 0.26 ± 0.14 cm).

**Figure 4:**
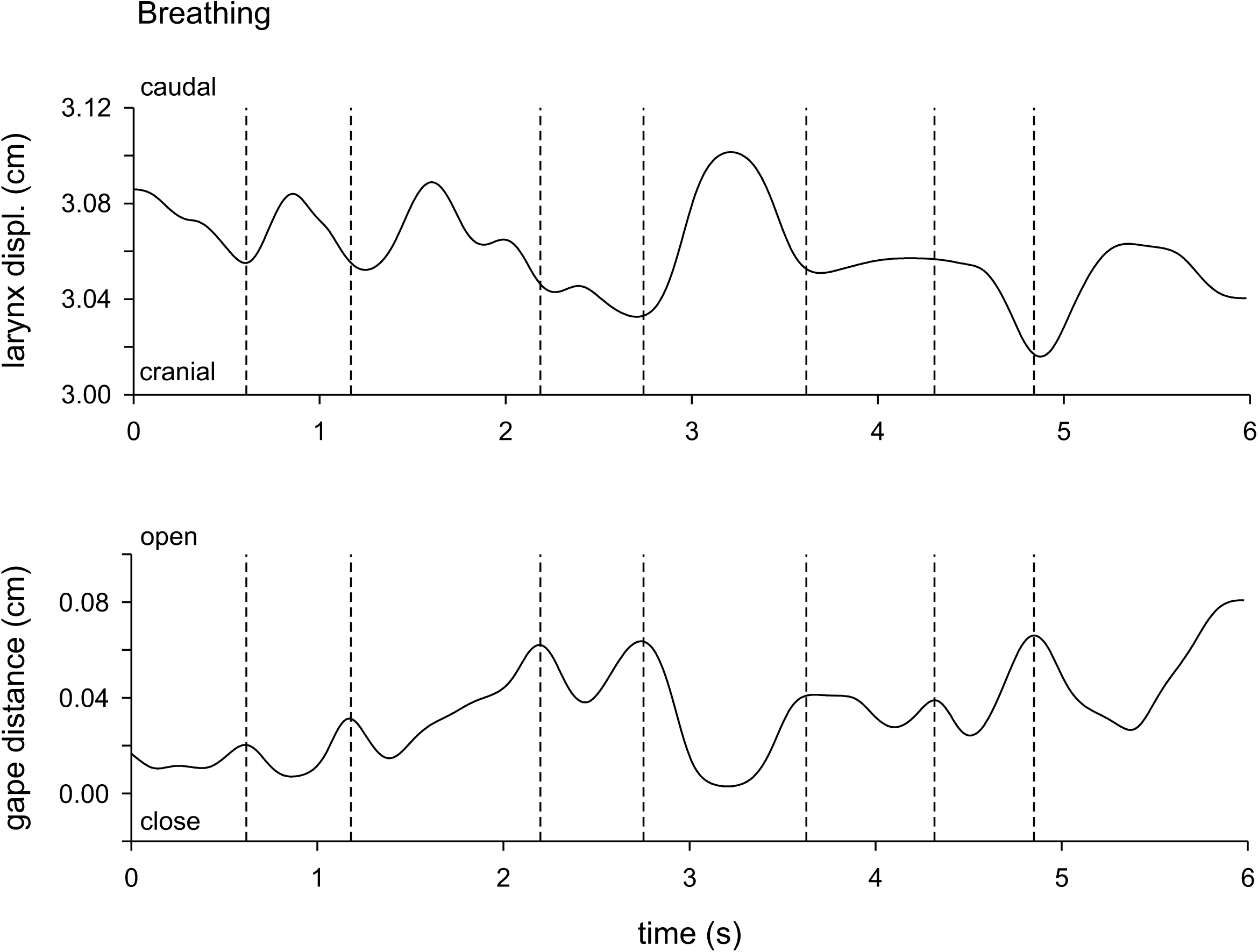
Laryngeal and lower jaw displacement profiles illustrating the coordination between jaw and anteroposterior larynx movements during breathing. The top graph illustrates the horizontal laryngeal displacement relative to the anterior lower jaw marker and the bottom graph illustrates the gape profile (jaw opening distance). Dashed lines indicate moments of maximal mouth opening which are often associated with maximal protraction of the larynx (lowest distance between the anterior jaw marker and the larynx). N = 2 animals.

**Figure 5:**
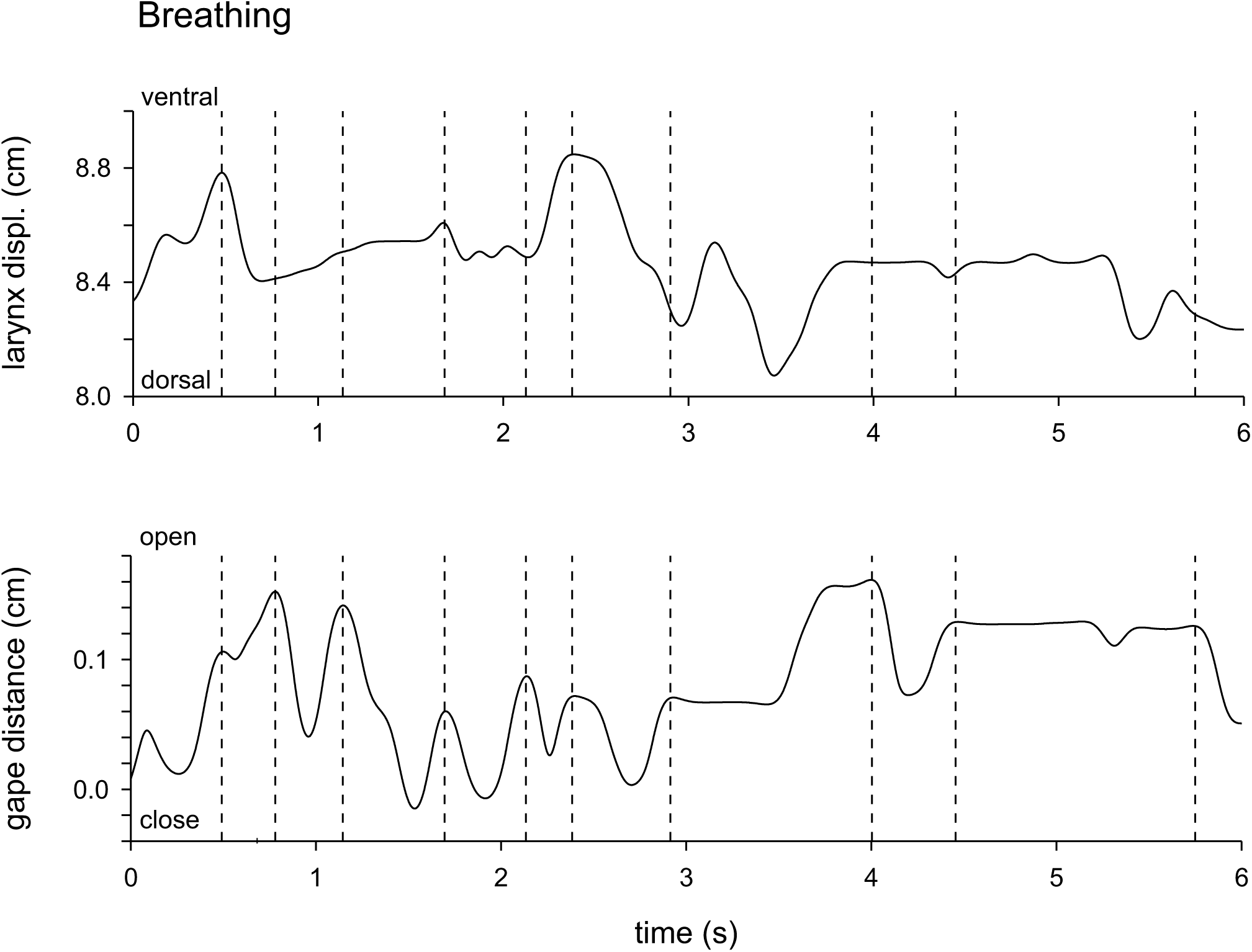
Laryngeal and lower jaw displacement profiles illustrating the coordination between jaw and dorsoventral larynx movements during breathing. The top graph illustrates the dorsoventral larynx displacement relative to the posterior upper jaw marker and the bottom graph illustrates the gape profile (jaw opening distance). Dashed lines indicate moments of maximal jaw opening. No clear synchronization between the dorso-ventral movements of the larynx and jaw opening are apparent. N = 2 animals

#### Vocalization

During vocalization, the movement of the larynx relative to the jaw is the reverse of that seen in breathing, with the larynx being pulled caudad and the distance between the tip of the lower jaw and the larynx increasing (Figs. 6, 7). Moreover, movements occur both in the horizontal and the vertical plane, different from breathing movements (Suppl. Videos 3-4). These movements are less regular and the posteroventral displacement of the larynx is generally associated with a slight increase in gape. However, in some instances gape increases before sound production and decreases during sound production, as illustrated in the last cycle in Fig. 6. Laryngeal movements are larger during vocalization and may encompass up to 5 mm in the horizontal plane (mean = 0.54 ± 0.40 cm) and 8 mm in the vertical plane (mean = 0.80 ± 0.71 cm). Jaw movements are also more pronounced and can be larger than 5 mm.

**Figure 6:**
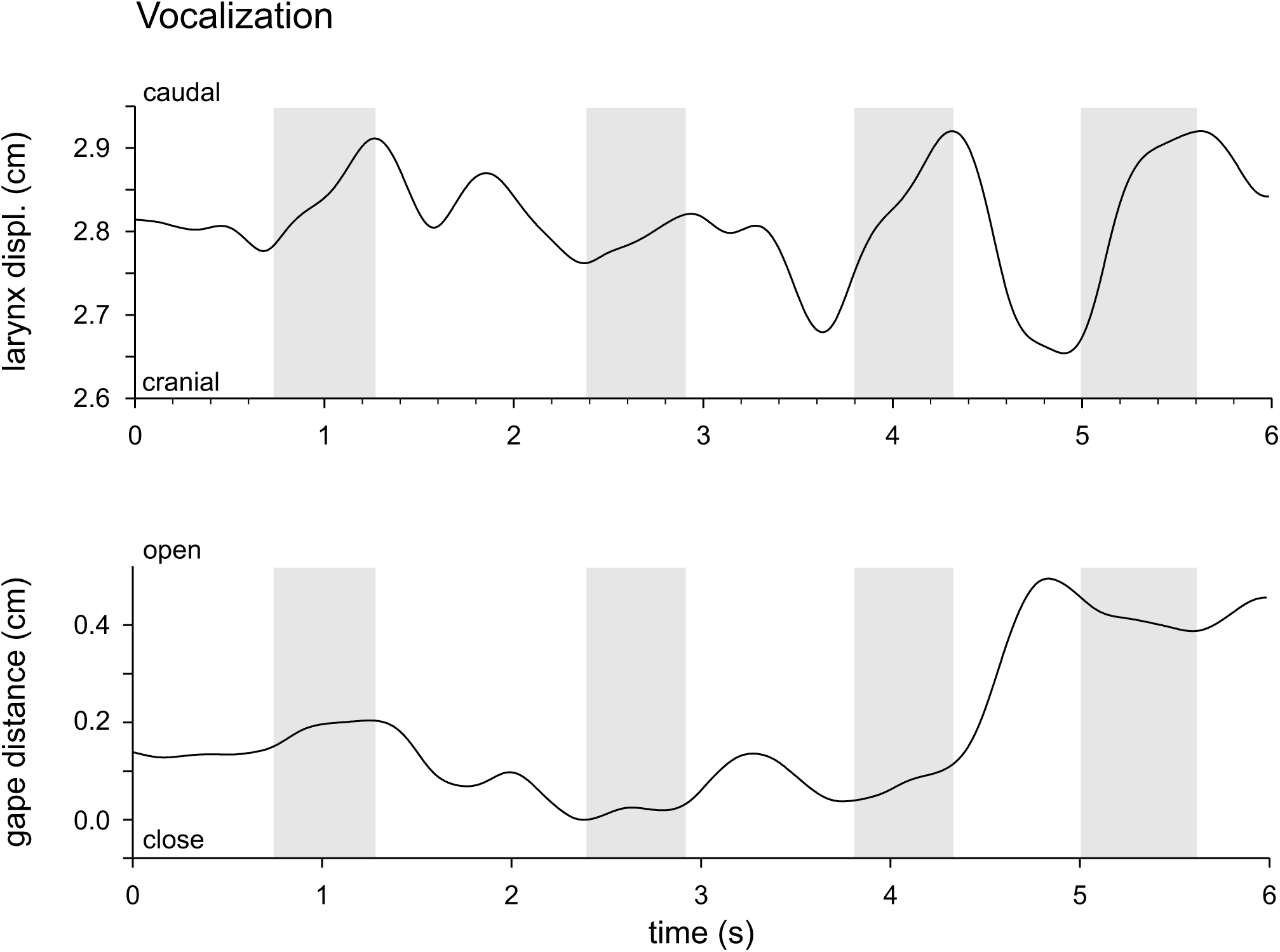
Displacement profiles of the larynx and lower jaws during vocalization. Grey areas denote times at which sound is produced by the gecko. During vocalization the larynx moves posteriorly with the posterior-most position being achieved towards the end of the vocalization. No clear coordination between vocalization and jaw opening is apparent, however. N = 2 animals

**Figure 7:**
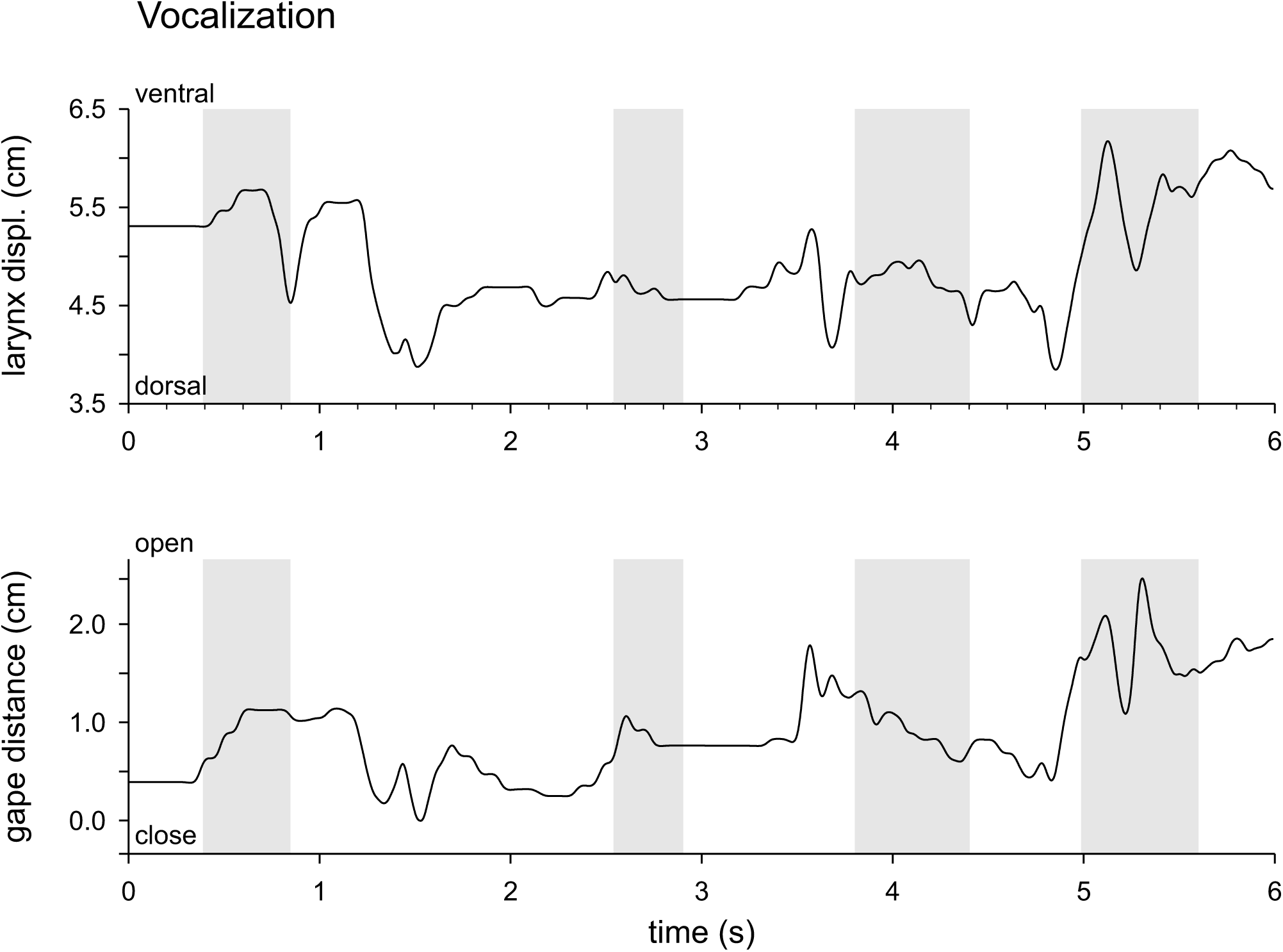
Displacement profiles of the larynx and the jaws during vocalization. Grey areas denote moments where sound is produced by the gecko. During vocalization no clear directionality of larynx movement is discernible can be observed and movements often include both ventral and dorsal displacements of the larynx. Similarly, no clear association between jaw opening and vocalization is observed.

#### Statistical analysis

The univariate analysis of variance testing for differences in laryngeal movements between breathing and vocalization was significant (antero-posterior displacement: *F*_1,29_ = 5.33; *P* < 0.001; dorso-ventral displacement: *F*_1,36_ = 6.34; *P* < 0.001) with geckos producing greater movements of the larynx during vocalization (antero-posterior: 0.54 ± 0.40 cm; dorso-ventral: 0.80 ± 0.71 cm) compared to breathing (antero-posterior: 0.06 ± 0.03 cm; dorso-ventral: 0.26 ± 0.14 cm). The cycle duration of the larynx movement was also significantly different between vocalization and breathing (antero-posterior displacement: *F*_1,29_ = 7.63; *P* = 0.01; dorso-ventral displacement: *F*_1,36_ = 8.55; *P* = 0.006) with cycles being shorter during vocalization (antero-posterior: 0.89 ± 0.31s; dorso-ventral: 0.70 ± 0.36 s) compared to breathing (1.28 ± 0.38 s; dorso-ventral: 0.93 ± 0.18 s). On average, the R^2^ value indicating the strength of the association between jaw and laryngeal movements during breathing was greater than during vocalization for the dorsal view sequences (breathing: 0.30 ± 0.23; vocalization: 0.14 ± 0.19) but not the lateral view sequences, where the pattern was opposite (breathing: 0.26 ± 0.40; vocalization: 0.50 ± 0.32). However, the statistical analysis suggests that the two types of behaviors were not significantly different in their coordination (dorsal view: *F*_1,11_ = 1.80; *P* = 0.21; lateral view: *F*_1,11_ = 1.40; *P* = 0.26).

### Anatomy of laryngeal motoneurons

After retrograde tracing of the glossopharyngeal nerve, we found labeled motoneurons exclusively located ipsilaterally within the hindbrain-spinal cord transition. Neurons were located at the level of the central canal or slightly more dorsally along the dorsoventral axis, in a central position along the dorso-ventral axis (Fig. 8 A). Axons could be followed from the nucleus ventrally (Fig. 8 B, arrow), where they exited the central nervous system as part of the glossopharyngeal nerve (N. IX). The shape of the individual neurons (N = 4 animals, n = 73 neurons) within the nucleus ranged from round to oval and fusiform (Fig. 8 B-D), with a mean minimal soma diameter of 18.1 µm (SEM: 1.1 µm; range = 5.4 – 48.8 µm), mean maximal soma diameter of 33.2 µm (SEM 1.6 µm; range = 12.8 – 74.1 µm) and a mean surface area of 646.2 µm^2^ (SEM: 50.2 µm^2^ ; range = 209.5 – 2202.2 µm^2^). The histograms depicting the minimal soma diameter, maximal soma diameter, and surface area show a left-skewed distribution (Fig. 8 E-G).

**Figure 8:**
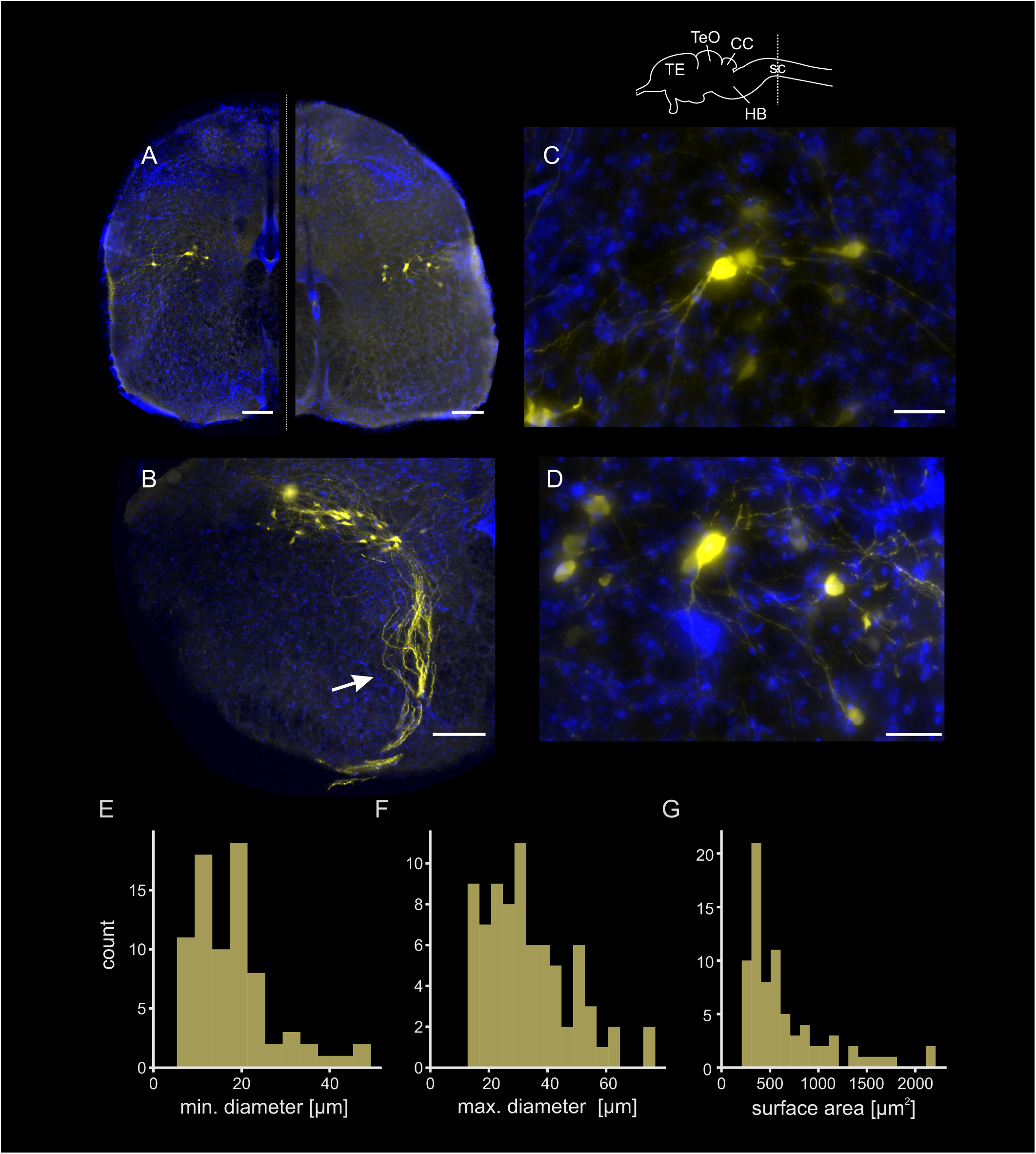
Anatomy of laryngeal motoneurons. Top: Sketch of the Gekko gecko brain. A) left: overview image of motoneurons within a transverse spinal cord section, right: overview of another section with backfilled motoneurons. B) Laryngeal motoneurons with axons (arrow) protruding towards the ventral horn of the spinal cord. C, D) Magnified image of motoneurons depicted in A (C: left side; D: right side). E) Histogram of min. soma diameter, bin size = 5 µm. F) Histogram of max. soma diameter, bin size = 5 µm. F) Histogram of surface area, bin size = 100 µm^2^. For Histograms in E-G: N = 4 animals/tracings, n = 73 neurons. For fluorescent images in A-D D: blue = DAPI, yellow = neurobiotin + streptavidin 546. Scalebars: A, B = 200 µm; C, D = 50 µm. Abbreviations: TE: Telencephalon; TeO: Optic tectum; CC: Cerebellum; HB: Hindbrain; sc: Spinal cord

## Discussion

### Kinematics of movement during breathing and sound production

The only prior study of the movements of the gecko larynx and glottis during sound production is that of Paulsen, in which direct observations of handheld Tokay geckos were employed (Paulsen, 1967). Paulsen stated that during the production of sounds, the larynx is pulled anteriorly and rotated to a more upright position, remaining in this orientation for several seconds before being retracted and returning to its resting position. Using an *in vitro* preparation, in which air was driven through the trachea, Paulsen then filmed the movements of the vocal cords at 8000 frames per second and showed that these move at a frequency between 300 and 400 Hz. Our results obtained from voluntarily emitted alarm calls were different: although we were unable to directly observe the movement of the glottis, the marker positioned next to the larynx allowed us to infer its movements. First, our results showed that movements of the larynx during sound production are significantly different from movements observed during breathing. Whereas during breathing, movements are mostly in the vertical plane (dorso-ventral movements), during vocalization, the glottis moved to a more posterior and ventral position, rather than to a more anterior and dorsal position, as suggested by Paulsen (Paulsen, 1967). Moreover, movements of the larynx during sound production are faster when compared to those when breathing and are of 600 and 700 ms in cycle duration, considerably shorter than what was observed by Paulsen (Paulsen, 1967). The combination of cineradiography and sound recordings allowed us to determine the onset of sound production and link it to the movements of the larynx, an advantage over the visual observations conducted by Paulsen. Interestingly, movements of the larynx were, on average, not coordinated with jaw movements, even though in some cases strong correlations were observed. This suggests that Tokay geckos rely strongly on movements of the larynx, which may be either coupled to or decoupled from movements of the jaws. These complex coordination patterns may be what allows them to produce different types of alarm calls as described here, but this remains to be investigated further.

### Neural networks controlling vocal behavior in non-avian reptiles

The neural control of sound production in non-avian reptiles is virtually unknown. A pair of exploratory studies conducted on *Gekko gecko* support the assumption of a generally conserved vocal pathway organization (Kennedy, 1975; Kennedy, 1981). In those studies, Kennedy managed to label vocal-associated motoneurons and was able to elicit vocalization by electrical stimulation of the midbrain, reportedly the periaqueductal gray. His tracings revealed motoneurons localized exclusively in the ipsilateral nucleus ambiguous in the caudal part of the medulla and the rostral part of the spinal cord (Kennedy, 1981). Laryngeal motoneurons showed a medio-lateral orientation but were not densely packed within the nucleus. These findings are in accord with our tracing results (Fig. 8). While Kennedy’s study showed neurons to be round or fusiform in shape and to exhibit maximal soma diameters in the range from 12-15 (round) to 20-30 µm (fusiform), motoneurons in our neural backfills (while revealing similar shapes and dendritic extends) span a broader size range of 12.8-74.1 µm.

Due to their conserved location at the caudal hindbrain/spinal cord boundary across vertebrates, vocal motoneurons were first regarded as having originated from a single developmentally or evolutionarily conserved progenitor pool (Bass and Baker, 2008; Bass et al., 2008). This classical view was recently challenged by data showing that the highly conserved *Phox2b* gene (presumed to be expressed in all vertebrate hindbrain branchial motoneurons) is present in the vocal motoneurons of frogs, mice, and humans but not in fish or avian reptiles (Albersheim-Carter et al., 2016). Generally, vocal motoneuron axons exit the brain via the twelfth cranial nerve in songbirds and the occipital nerve (potential homolog of the N.XII) in fish, and via the ninth and tenth cranial nerves in frogs and mammals (Barkan and Zornik, 2020). While evolutionary considerations previously suggested convergent evolution (Barkan and Zornik, 2020) of vocal traits, recent evidence suggested that a common ancestor in choanate vertebrates was already vocal (Barkan and Zornik, 2020; Jorgewich-Cohen et al., 2022). Due to their rich vocal behavior, gekkotans, turtles and crocodilians offer a unique opportunity for improving our understanding of vocal pathway(s) and the control and modulation of sound features.

Our *in vitro* tracings further revealed that the gecko hindbrain remains stable in Ringer solution for multiple days at appropriate temperature. While we have not yet recorded from neurons in such preparations yet, these first results indicate that *in vitro* electrophysiological and anatomical studies, similar to the ones performed on diverse motor patterns, including locomotor activity, respiration, and vocal behavior (Kelley et al., 2017; Rhodes et al., 2007; Zornik and Kelley, 2008), are likely to be successful in *G. gecko* as well. If an *in vitro* whole brain preparation of *G. gecko* would allow for the production of fictive calls (i.e., action potential volleys with features specific to the *G. gecko* sounds in the nerves associated with the control of sound production), then such a preparation would greatly facilitate studying vocal networks irrespective of respiratory activation, as already shown for the amphibian *Xenopus laevis*. The ability of *G. gecko* to produce different sound types in different contexts might thus allow for the study of the neural substrates of context-dependent sound production on easily identifiable fictive motor patterns (i.e., nerve activity).

As non-avian reptiles have their sound production organ (i.e., the larynx) located at the cranial end of the trachea (in contrast to the syrinx of avian reptiles), this facilitates the study of its biomechanics and allows for more direct comparisons with mammalian sound production. Consequently, *Gekko gecko* could serve as a reference taxon for comparative studies of sound production as the reptile counterpart of the midshipman fish among teleosts (Bass and Remage-Healey, 2008; Chagnaud et al., 2021), the African clawed frog among amphibians, and the zebra finch (Brenowitz et al., 1997; Düring et al., 2013; Elemans, 2014) among birds.

## Acknowledgements

We acknowledge the help of Birgit Rönfeld in histological processing and of Renaud Boistel for lending audio equipment.

## Author contributions

**Conceptualization**: LK; **Software**: TJ, RG; **Validation**: RG, LK, TJ, DTO, BPC, AH; **Formal analysis**: RG, TJ, DTO, BPC, AH; **Investigation**: RG, LK, BPC, AH; **Resources**: BPC, AH; **Data Curation**: RG, LK, BPC, AH; **Writing – original draft preparation**: RG, TJ, BPC, AH; **Writing – review and editing**: RG, LK, TJ, BPC, AH; **Visualization**: RG, TJ, BPC, AH; **Supervision**: AH, BPC; **Project administration**: RG, LK, BPC, AH; **Funding acquisition:** RG, LK, BPC, AH;

## Data availability

Data will be made available upon request.

## List of abbreviations

BDA: biotinylated dextran amine
CC: cerebellum
DAPI: 4′,6-diamidino-2-phenylindole
FFT: Fast Fourier Transform
fps: frames per second
HB: hindbrain
MFCC: mel-frequency cepstral coefficients
MNHN: Muséum national d’Histoire naturelle
PB: phosphate buffer
PCA: principle component analysis
PFA: paraformaldehyde
PSD: power spectral density
sc: spinal cord
SVL: snout vent length
TE: telencephalon
TeO: optic tectum
Tx-100: Triton X-100

