## Supplemental Information for "*Gekko gecko* as a model organism for understanding aspects of laryngeal vocal evolution"

**Supplementary material**

**Supplementary table 1:**

| Individual | Total | Type 1 | Type 2 | Type 3 | Sex |
|---|---|---|---|---|---|
| **3** | 32 | 9 | 7 | 16 | male |
| **4** | 14 | 7 | 3 | 4 | male |
| **5** | 12 | 4 | 2 | 6 | male |
| **6** | 14 | 7 | 5 | 2 | male |
| **7** | 56 | 16 | 8 | 32 | male |
| **8** | 28 | 11 | 9 | 8 | female |
| **Sum** | **156** | **54** | **34** | **68** | |

**Supplementary table 2:**

| component of the study | number of animals used (N) | number of events observed (n) | number of events observed per individual | sex of animals |
|---|---|---|---|---|
| sound recordings | 6 | 156 | see supplementary table 1 | 5 males<br>1 female |
| PCA of sound recordings | 6 | 147 | see supplementary table 1 | 5 males<br>1 female |
| X-ray analysis (vocalization sequences) | 2 | 10 | 5 sequences per animal | 2 males |
| X-ray analysis (breathing sequences) | 2 | 2 | 2 sequences per animal | 2 males |
| absolute breathing events (lateral view) | 2 | 11 | animal 1 = 5 events<br>animal 2 = 6 events | 2 males |
| absolute vocalization events (lateral view) | 2 | 14 | animal 1 = 14 events<br>animal 2 = 6 events | 2 males |
| absolute breathing events (dorsal view) | 2 | 13 | animal 1 = 8 events<br>animal 2 = 5 events | 2 males |
| absolute vocalization events (dorsal view) | 2 | 25 | animal 1 = 8 events<br>animal 2 = 17 events | 2 males |
| neuronal backfills | 4 | 4 | 1 backfill per animal | 4 males |

**Supplementary figure 1:**

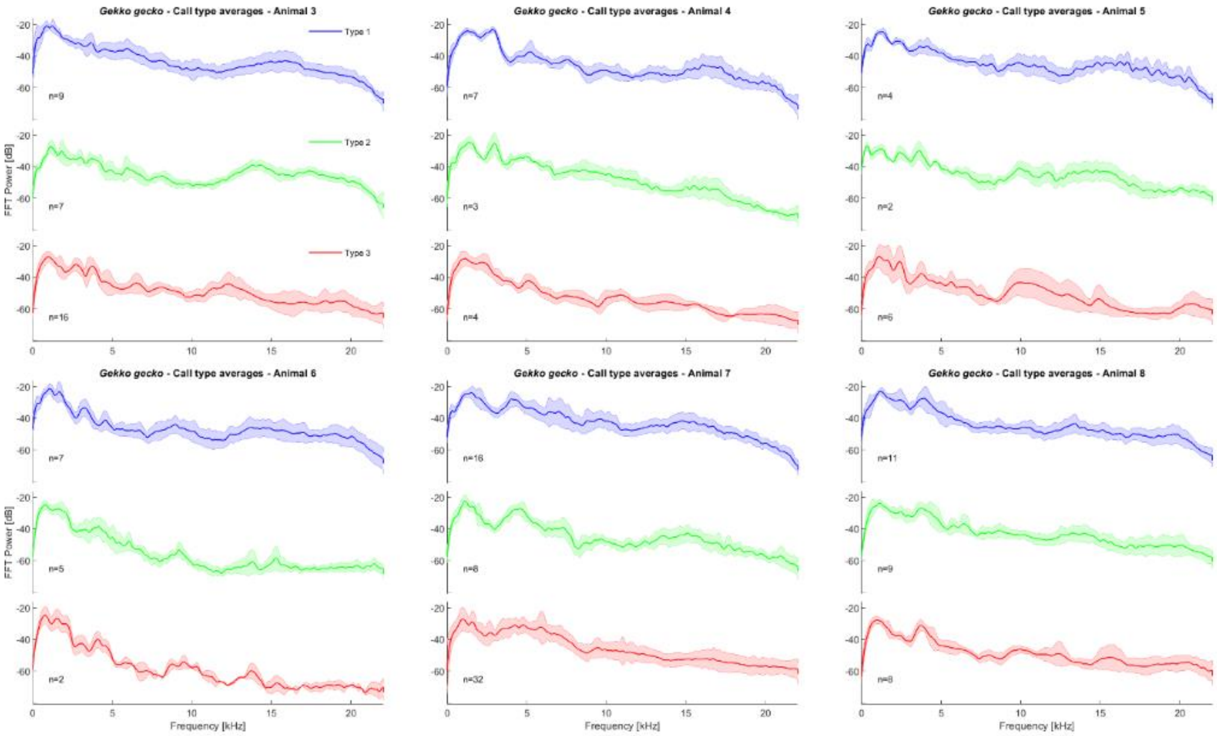

**Supplementary figure 2:**

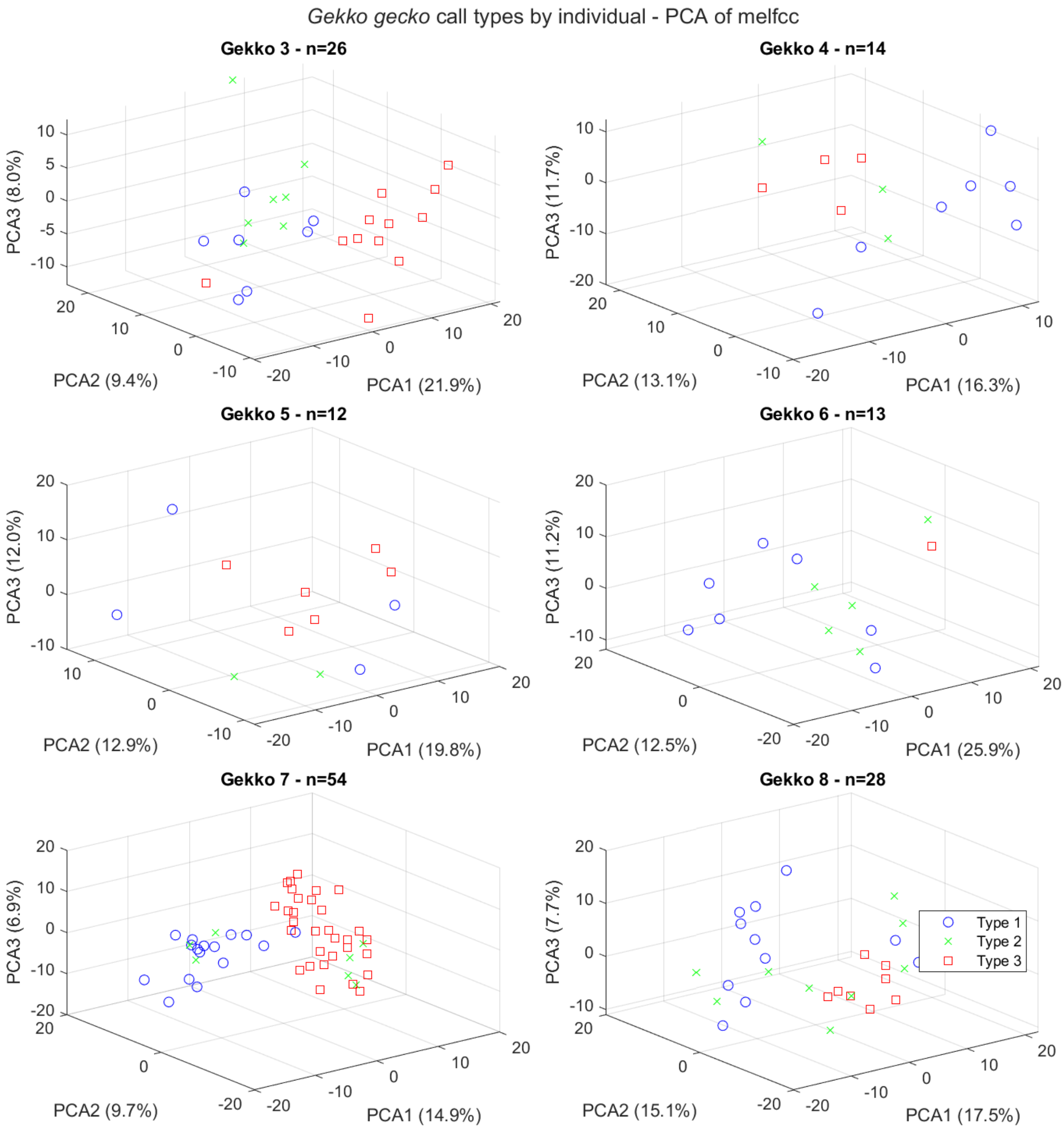

Supplementary figure 3:

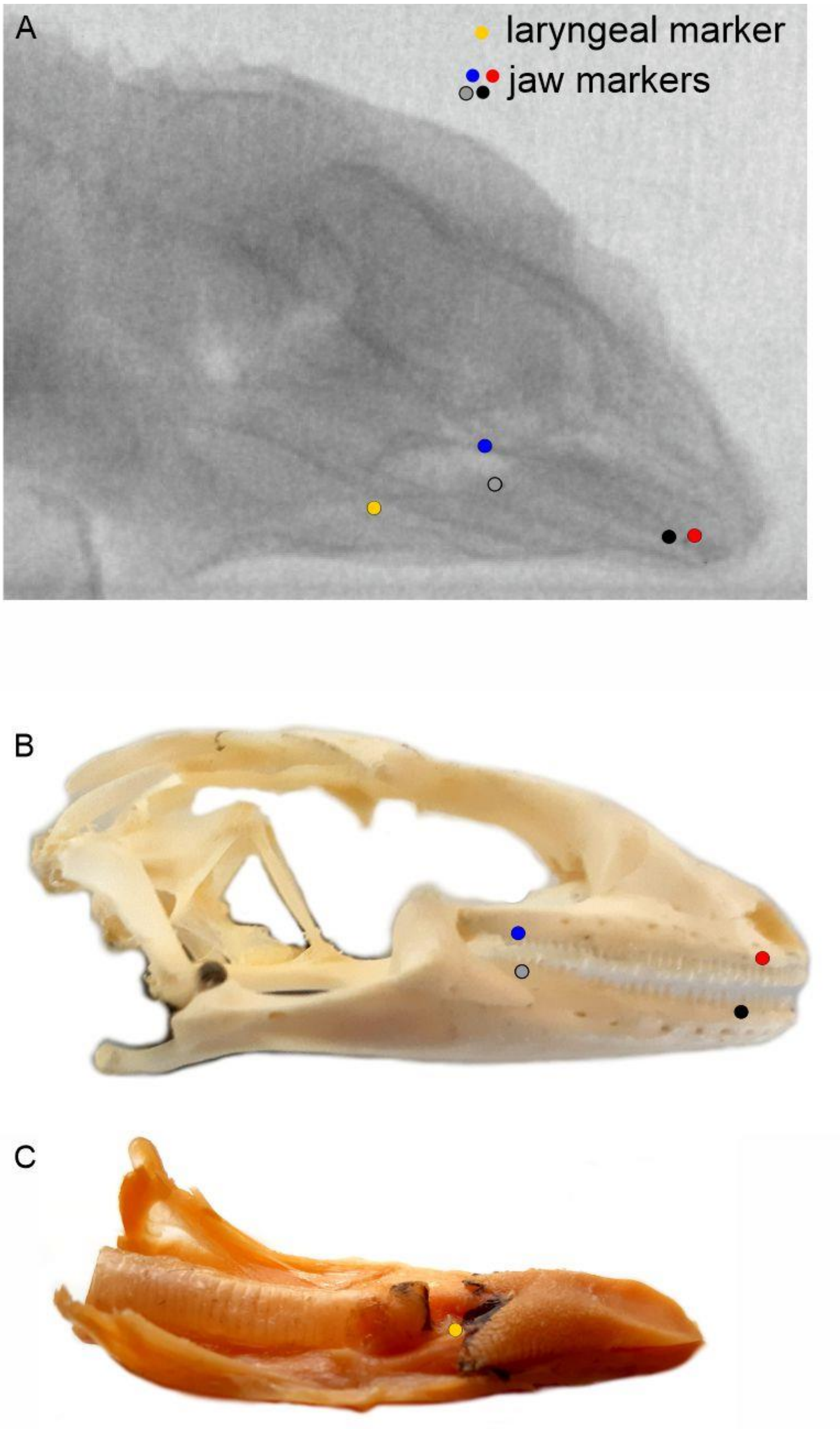

**Supplementary legends**

**Supplementary table 1:** Distribution of number of call types produced by 6 *Gekko gecko* individuals.

**Supplementary table 2:** Sample sizes used in different components of the study.

**Supplementary figure 1:** Mean frequency spectra of the three call types produced by each individual animal. Solid lines show averages and shaded areas depict ± 1 standard deviation.

**Supplementary figure 2:** Results of PCA on the MFCCs of vocalizations for each individual *Gekko gecko*.

**Supplementary figure 3:** (A) X-Ray image of a *Gekko gecko* individual, showing the implanted markers. (B) Schematic of a *Gekko gecko* skull as a reference with location of the implanted markers as an overlay. (C) Schematic of a *Gekko gecko* tongue with implanted laryngeal marker position as an overlay. Colors: blue, red, gray and black = skull markers, yellow = laryngeal marker.

**Supplementary movie 1:** Displacement of jaw and laryngeal markers visualized by dorso-ventral cineradiography of a *Gekko gecko* during breathing.

**Supplementary movie 2:** Displacement of jaw and laryngeal markers visualized by lateral cineradiography of a *Gekko gecko* during breathing.

**Supplementary movie 3:** Displacement of jaw and laryngeal markers visualized by dorso-ventral cineradiography of a *Gekko gecko* during vocalization.

**Supplementary movie 4:** Displacement of jaw and laryngeal markers visualized by lateral cineradiography of a *Gekko gecko* during vocalization.
